# Distinguishing Polymer Turnover from *De Novo* Biosynthesis Reveals Dynamic Cell Wall Remodeling During *Aspergillus fumigatus* Conidial Germination

**DOI:** 10.64898/2026.07.31.740354

**Authors:** Yohara K. Ranasinghe, Ankur Ankur, Jean-Paul Latgé, Tuo Wang

**Affiliations:** Department of Chemistry, Michigan State University, East Lansing, MI, USA; Unité des Aspergillus, Institut Pasteur, Paris, France

**Keywords:** Fungi, Pathogen, Carbohydrate, *Aspergillus*, Solid-state NMR, Cell wall, Germination, Conidia, Glucan, Chitin

## Abstract

Dynamic remodeling of extracellular matrices underlies development, environmental adaptation, and host-pathogen interactions, yet distinguishing polymer turnover from *de novo* biosynthesis in intact cells remains a major challenge. Here, we combined high-resolution solid-state NMR with selective ^13^C-labeling strategies to distinguish pre-existing cell-wall polymers from newly synthesized polysaccharides during *Aspergillus fumigatus* conidial germination. Germination was accompanied by substantial remodeling of the rigid cell wall, characterized by decreased β-1,3-glucan and increased chitin and α-1,3-glucan, whereas the mobile wall fraction remained comparatively stable except for the emergence of galactosaminogalactan. Surprisingly, β-1,3-glucan turnover proceeded independently of the major β-1,3-glucanases encoded by the *A. fumigatus* genome and was dispensable for germination. Instead, isotope-labeling experiments revealed that newly assimilated carbon is preferentially directed toward α-1,3-glucan biosynthesis, whereas deletion of the α-1,3-glucan synthase genes triggered compensatory accumulation of chitin and β-1,3-glucan. These results reveal a compartmentalized cell-wall remodeling program that coordinates selective turnover with *de novo* polysaccharide synthesis during fungal germination and establish isotope-edited solid-state NMR as a general approach for distinguishing inherited from newly synthesized polymers in complex carbohydrate matrices.

## INTRODUCTION

Severe fungal diseases impose a major global health and economic burden, contributing to millions of deaths each year and substantial treatment costs worldwide^[1–2]^. Among the fungal pathogens responsible for invasive disease, *Aspergillus fumigatus* is one of the most clinically significant, particularly because of its ability to cause life-threatening infections in immunocompromised individuals^[3–5]^ as well as allergic reactions in immunocompetent individuals^[6]^. This fungus produces tiny airborne spores, commonly known as conidia, that can easily travel through the respiratory system.

The onset of infection relies on germination of the conidia, a process involving two major morphological changes: isotropic swelling and polarized growth, the latter of which leads to formation of a filamentous network that can invade the lung. Despite the central role of this morphological transition to disease progression early germination phase of *A. fumigatus* remains incompletely understood. A major barrier to resolving this stage is the complexity and rapid remodeling of the fungal cell wall^[7–8]^, which simultaneously provides adaptive mechanical strength and serves as an armor between the fungus and the host immune system^[9–10]^. It comprises a polysaccharide-rich scaffold arranged in an inner rigid core and outer mobile domain and enables dynamic responses to environmental cues, including commitment to germination^[11–12]^.

Recent studies have begun to resolve germination-associated cell wall remodeling at the molecular level, with imaging and biochemical approaches revealing stage-specific changes in conidial surface architecture and enzyme activity during germination^[13–18]^. However, it remains unclear whether the cell wall changes accompanying germination are driven primarily by the remodeling of pre-existing conidial wall material or/and by *de novo* synthesis from newly acquired nutrients. Because cell wall remodeling occurs within a structurally heterogeneous matrix composed of structurally diverse polysaccharide domains, resolving these contributions requires molecular-level methods capable of monitoring intact cell walls *in situ*. In this study, we combine high-resolution solid-state NMR spectroscopy^[19-21]^ with a series of ^13^C-labeling strategies to distinguish pre-existing conidial biomass from newly synthesized carbohydrates during *A. fumigatus* germination. This approach allows us to track polysaccharide turnover, identify major biosynthetic and remodeling pathways, and define how the fungal cell wall is reorganized during the transition from dormant conidia to actively growing germlings.

## RESULTS

### Germination remodels the pre-existing conidial cell wall architecture

Early stages of germination are characterized by pronounced morphological transitions, beginning with conidial swelling and progressing to polarized growth. Light microscopy images of *A. fumigatus* captured resting conidia at 0 h, swollen conidia at 5 h, and germinating conidia exhibiting germ tube emergence at 7 h (**Figure 1A**). The appearance of germ tubes at 7 h marks the transition from isotropic expansion to polarized hyphal growth. Quantification of cell populations in triplicates revealed a progressive decline in the proportion of resting conidia, accompanied by a corresponding increase in swollen and germinating cells over time (**Figure 1B**). At 5 h, most cells had entered the isotropic growth phase, with approximately 80% of the population becoming swollen, whereas by 7 h, approximately 70% of conidia had formed germ tubes, demonstrating a coordinated transition to polarized growth under the conditions examined.

**Figure 1.**
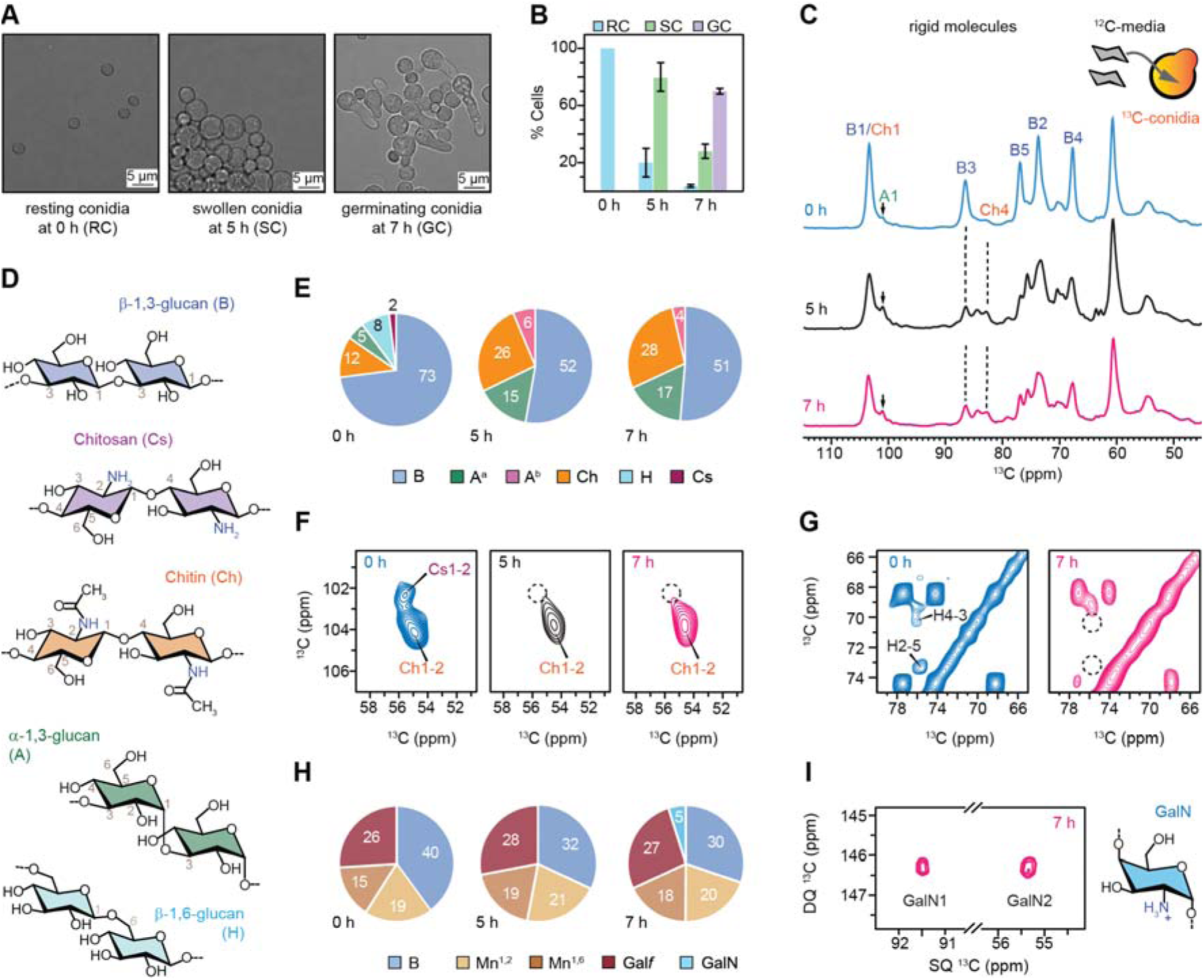
Remodeling of cell-wall polysaccharides during *A. fumigatus* conidial germination. (**A**) Representative brightfield micrographs showing morphological transition of *A. fumigatus* from resting conidia at 0 h to swollen conidia at 5 h and germinating conidia at 7 h. (**B**) Average cell populations at each germination stage show the progressive loss of resting conidia and accumulation of swollen and germinating cells over time. Error bars represent s.d. across triplicates. (**C**) 1D ^13^C CP spectra detecting rigid molecules derived from the pre-existing conidial carbon biomass in ^13^C-conidia grown in unlabeled media (inset). Major polysaccharide signals are assigned abbreviations as β-1,3-glucan (B), α-1,3-glucan (A), chitosan (Cs), chitin (Ch), and β-1,6-glucan (H). Dashed lines highlight changes in signature peaks of chitin and β-1,3-glucan. (**D**) Representative chemical structures and abbreviations for the major rigid cell-wall polysaccharides. Key carbon sites are numbered. (**E**) Molar composition of rigid polysaccharides based on peak intensity analysis of 2D ^13^C CORD spectra. (**F**) Zoomed regions of 2D CORD spectra show the presence of chitosan at 0 h and absence at later stages. (**G**) Zoomed region of 2D CORD spectra show changes in β-1,6-glucan-associated signals between 0 h and 7 h. (**H**) Molar composition of mobile polysaccharides calculated using peak intensity of 2D ^13^C DP refocused J-INADEQUATE spectra. (**I**) Zoomed region of J-INADEQUATE spectra showing galactosamine signals absent in 0 h but detected at 7 h, neosynthesized using carbons already available in the conidia.

To track the possible reuse of pre-existing cell wall polysaccharides during germination, we transferred uniformly ^13^C-labeled conidia into unlabeled medium. In this labeling scheme, molecules newly synthesized from the unlabeled medium are not detected, allowing us to unambiguously monitor the fate and reuse of pre-existing ^13^C-labeled molecules already present in the cell. In dormant conidia at 0 h, the rigid domain of *A. fumigatus* cell wall was dominated by β-1,3-glucan, as shown by its carbon-3 (B3) peak at 87 ppm, with minor contributions from α-1,3-glucan, identified by the carbon-1 (A1) peak at 101 ppm, and chitin, identified by the carbon-4 (Ch4) peak at 83.5 ppm (**Figure 1C, D**). β-1,3-glucan accounted for approximately 73% of the rigid carbohydrates (**Figure 1E**). At the swollen stage (5 h), the relative contribution of β-1,3-glucan decreased to approximately 52%, whereas chitin and α-glucan increased to 26% and 15%, respectively (**Figure 1C, E**). This spectral profile and overall cell wall composition were largely maintained at the germinating stage (7 h). These results show that the transition from dormant to swollen conidia involves substantial remodeling of pre-existing rigid polysaccharides, converting some β-1,3-glucans to chitin and α-1,3-glucan, whereas subsequent germ tube emergence relies predominantly on *de novo* cell wall synthesis with minimal reutilization of existing rigid polysaccharides.

High-resolution from 2D ^13^C-^13^C CORD spectra acquired using ^1^H-^13^C cross polarization (CP) further resolved the selective removal of pre-existing chitosan and β-1,6-glucan during germination. Chitosan was unambiguously detected in dormant conidia by its characteristic C1-C2 (Cs1-2) cross-peak at (102.8, 56.7) ppm, which disappeared completely in swollen and germinating conidia (**Figure 1F**; **Fig. S1**). Similarly, β-1,6-glucan, identified by its characteristic H4-3 and H2-5 cross-peaks at (70.8, 76.8), and (74.2, 75.8) ppm, respectively (**Figure 1G**). Because germination was performed in unlabeled medium, the loss of chitosan and β-1,6-glucan signals from pre-labeled conidia indicates degradation or chemical modification of the original dormant conidial wall architecture. However, it should be noted that solid-state NMR cannot distinguish a heterogeneous polymer containing both N-acetylglucosamine (GlcNAc) and glucosamine (GlcN) residues from a mixture of separate GlcNAc and GlcN homopolymers. Therefore, the observed chitosan signals may alternatively reflect glucosamine residues incorporated within a predominantly GlcNAc polymer rather than a distinct GlcN polymer.

Mobile polysaccharides were characterized using a 2D refocused J-INADEQUATE experiment with ^13^C direct polarization (DP) and a short recycle delay, which selectively detect signals from mobile components with rapid ^13^C spin-lattice relaxation. Under the same labeling scheme, the mobile domain of dormant conidia was dominated by β-1,3-glucan and galactomannan (GM), including α-1,2-mannose, α-1,6-mannose, and galactofuranose (Gal*f*) residues (**Figure 1H**; **Fig. S1**). Similar to the rigid fraction, redistribution of pre-existing carbon in the mobile fraction occurred primarily during the transition from dormant to swollen conidia, as reflected by a decrease in β-1,3-glucan signals and a concomitant increase in galactomannan signals (**Figure 1H**). In contrast, a unique feature of the mobile fraction was the appearance of GAG-associated GalN resonances at 7 h (**Figure 1I**), indicating the onset of GAG deposition during early morphogenesis and the production of surface-associated polysaccharides as conidia transitioned to polarized germ tube growth.

### Partial **β**-1,3-glucan degradation does not significantly affect conidial germination

As linear β-1,3-glucan chains are synthesized by the glucan synthase complex and extruded across the plasma membrane into the periplasmic space, remodeling of the pre-existing cell wall is required to facilitate their incorporation^[22–23]^. One biochemical process associated with germination is the localized softening of the cell wall by glycosyl hydrolases, which reduces wall rigidity and enables cell expansion. Upon reaching the cell wall matrix, newly synthesized β-1,3-glucan undergoes extensive structural remodeling, including branching, elongation, and controlled degradation, before being integrated into the existing wall architecture. These remodeling events are essential for morphogenetic processes such as conidial swelling, germ tube emergence, and lateral hyphal formation, all of which require localized weakening and reorganization of the cell wall to accommodate the growth of new cellular structures. β-1,3-glucan-hydrolyzing enzymes involved in this process are broadly classified into endo-β-1,3-glucanases and exo-β-1,3-glucanases^[17]^. Endo-β-1,3-glucanases cleave internal glycosidic linkages within glucan chains in a relatively random manner, whereas exo-β-1,3-glucanases sequentially release glucose residues from the non-reducing ends of the polymers (**Figure 2A**).

**Figure 2.**
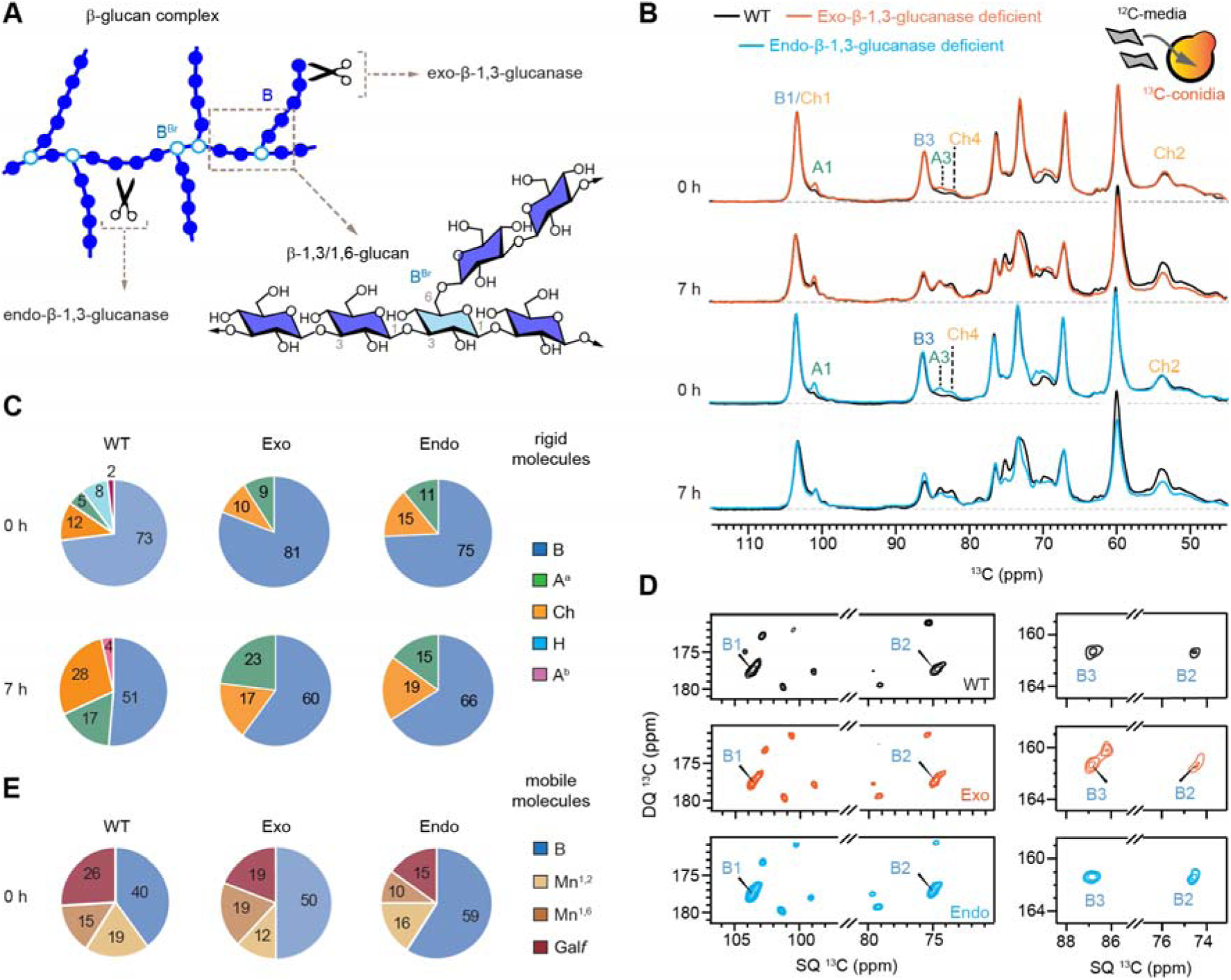
β-1,3-glucanase deficiency alters remodeling of rigid conidial wall during germination. (**A**) Schematic representation of β-glucan remodeling by endo- and exo-β-1,3-glucanases. (**B**) 1D ^13^C CP spectra of wild-type (black), exo-β-1,3-glucanase-deficient (orange), and endo-β-1,3-glucanase-deficient (blue) strains prepared by growing ^13^C-labeled conidia in unlabeled media for 0 h and 7 h. Horizontal dashed line (grey) shows spectral baseline. (**C**) Molar composition of rigid polysaccharides of wild-type (left), exo-β-1,3-glucanase-deficient (middle), and endo-β-1,3-glucanase-deficient (right) grown for 0 h (top) and 7 h (bottom) estimated from the peak volumes of resolved peaks of 2D ^13^C CORD spectra. (**D**) Selected regions of 2D ^13^C refocused J-INADEQUATE spectra of 0-h samples show differing intensities of β-1,3-glucan peaks in resting conidia across strains. (**E**) Molar composition of the mobile molecules in the 0 h conidia of these three strains.

To understand whether the decrease in rigid β-1,3-glucan observed during conidial germination depends on β-1,3-glucanase-mediated cell wall remodeling, we analyzed two glucanase-deficient strains^[24–25]^, which were constructed from the deletion of the major glucanase genes identified in the *A. fumigatus* genome belonging to all members of the GH16, 55 and 81 family from the CAZy database^[26]^. These mutants enabled us to investigate the relative contributions of exo- and endo-β-1,3-glucanase activities to the remodeling of pre-existing conidial cell wall polysaccharides during germination. 1D ^13^C CP spectra revealed that β-1,3-glucan remained the predominant rigid carbohydrate component in both mutant strains at 0 h and 7 h (**Figure 2B**). At 0 h, strong β-1,3-glucan signals, including the characteristic B1 and B3 peaks, were observed in both mutants and were comparable to those of the wildtype strain. Furthermore, germination proceeded with similar kinetics across all strains, indicating that deletion of the examined β-1,3-glucanases does not substantially impair the germination process under the conditions tested.

Compositional analysis revealed similar initial cell wall carbohydrate profiles among the wild-type and glucanase-deficient strains, with β-1,3-glucan accounting for 73-81% of the rigid carbohydrate pool at 0 h, indicating that disruption of either glucanase had negligible effect on the overall abundance of rigid β-1,3-glucan in resting conidia (**Figure 2C**). During germination, the relative contribution of rigid β-1,3-glucan decreased in all three strains, but the magnitude of this decrease differed: β-1,3-glucan was reduced by 21-22% in the wildtype and exo-β-1,3-glucanase-deficient strains, whereas the endo-β-1,3-glucanase-deficient strain exhibited a markedly smaller reduction by only 9% (**Figure 2C**).

These results indicate that deletion of the tested exo-β-1,3-glucanases does not prevent the germination-associated reduction in rigid β-1,3-glucan, suggesting that exo-glucanase activity is not a major determinant of this remodeling process. In contrast, loss of the examined endo-β-1,3-glucanases attenuated the decrease in rigid β-1,3-glucan.

The reduction in rigid β-1,3-glucan during germination was accompanied by concomitant changes in other cell wall polysaccharides, revealing distinct remodeling trajectories among the strains. In the wild type, chitin increased from 12% at 0 h to 28% at 7 h, while α-1,3-glucan increased from 5% to 17% (**Figure 2C**), indicating extensive restructuring of the rigid wall matrix as β-1,3-glucan content declined. A similar pattern was observed in the exo-β-1,3-glucanase-deficient strain, where chitin increased from 10% to 17% and α-1,3-glucan reached 9 to 23% at 7 h. The endo-β-1,3-glucanase-deficient strain exhibited a modest increase in chitin (15% to 19%) and α-1,3-glucan (11 to 15%), paralleling its reduced loss of β-1,3-glucan. Loss of glucanase activity does not substantially impair chitin and α-1,3-glucan remodeling and may promote compensatory α-1,3-glucan accumulation paralleling its reduced loss of β-1,3-glucan.

In both exo- and endo-β-1,3-glucanase-deficient strains, β-1,6-glucan signals were not detected in both resting and germinating conidia, whereas these characteristic correlations were readily observed in resting conidia of the wild-type strain before disappearing during swelling and germination (**Figs. S1**, **S2**). The reduction in β-1,3-glucan during conidial germination was approximately 30% in the parental strain and 12-26% in the exo- and endo-β-1,3-glucanase-deficient strains. Despite these differences in the extent of β-1,3-glucan degradation, no significant differences in conidial germination were observed among the strains.

In addition, the dormant conidia of both glucanase-deficient strains exhibited a substantial enrichment of the mobile β-1,3-glucan pool, which accounted for 50% and 59% of the mobile carbohydrates in the exo- and endo-β-1,3-glucanase-deficient strains, respectively, compared with a lower abundance in the wildtype strain (**Figure 2D, E**). The enrichment of mobile β-1,3-glucan in dormant conidia of both glucanase-deficient strains suggests that in the absence of these enzymes, incomplete glucan remodeling likely results in the accumulation of more mobile β-1,3-glucan populations, indicating that β-1,3-glucanases influence cell wall architecture prior to germination.

All these results indicate that even modest changes in the rigid and mobile cell wall polysaccharides can lead to, or be associated with, changes in other cell wall polymers, but these moderate changes did not significantly alter the percentage of conidial germination.

### α-1,3-glucan turnover and synthesis drive cell wall remodeling during germination

To comprehensively track the kinetics of cell wall remodeling and distinguish *de novo* synthesis from the reutilization of pre-existing cell wall components, we employed three complementary ^13^C-labeling strategies (**Figure 3A**): (i) uniformly ^13^C-labeled conidia germinated in unlabeled medium to monitor the fate and reuse of pre-existing carbon pools, as described above; (ii) unlabeled conidia germinated in ^13^C-labeled medium to selectively detect newly synthesized molecules; and (iii) uniformly ^13^C-labeled conidia germinated in ^13^C-labeled medium to provide a comprehensive profile of total cell wall polysaccharides during germination for comparison with the selective labeling experiments.

**Figure 3.**
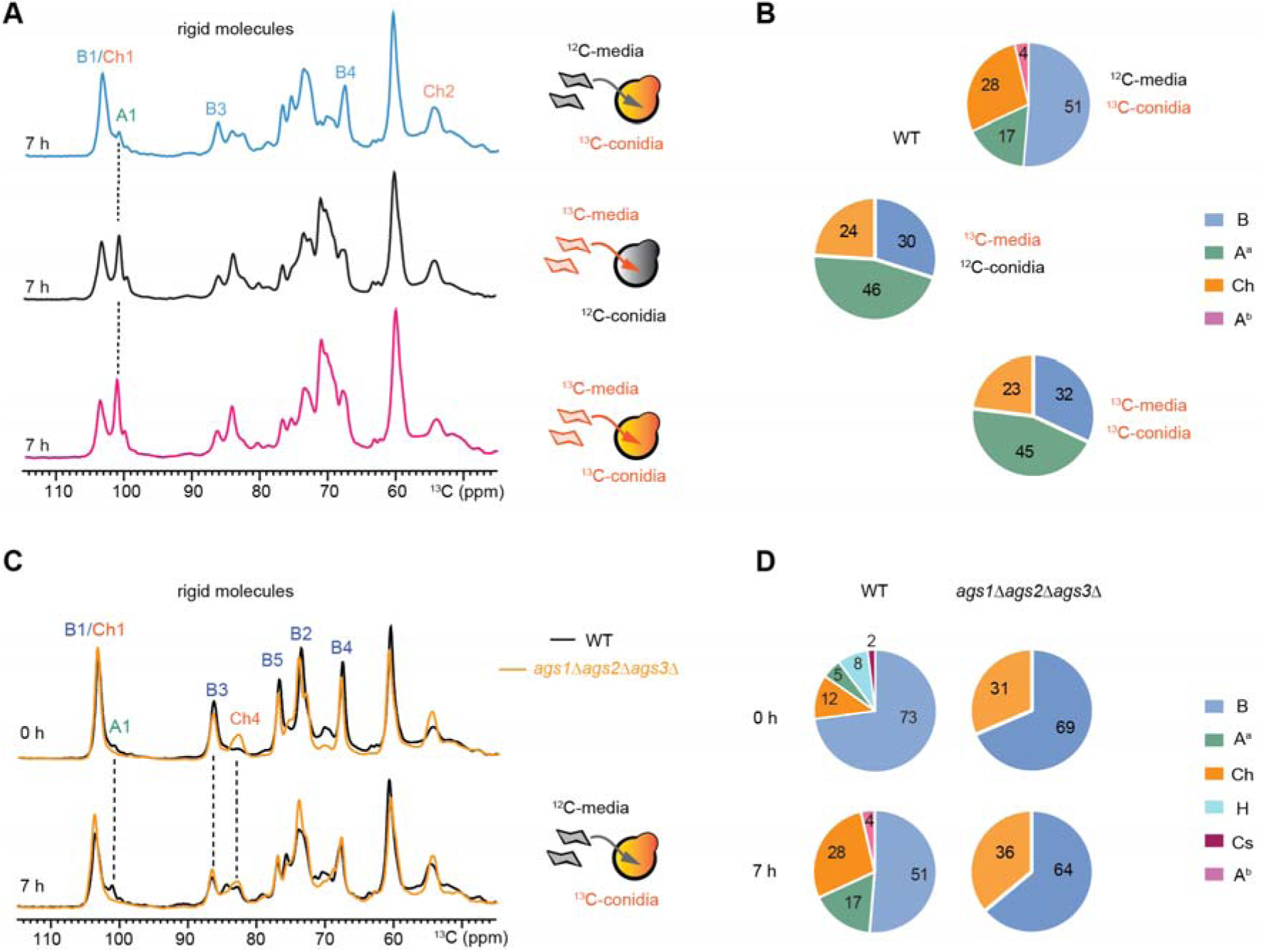
Neo-synthesized α-glucans contribute to remodeling rigid walls during germination. (**A**) 1D ^13^C CP spectra showing rigid cell wall polysaccharides of 7 h germinating conidia labeled using different isotopic enrichment schemes. The scheme on the right side illustrates the corresponding ^13^C labeling strategy for the corresponding sample measured in each spectrum. (**B**) Molar composition of polysaccharides in the rigid cell wall domain of samples grown to 7 h, calculated from the peak volumes from 2D ^13^C CORD spectra corresponding to the 1D spectra in panel A. (**C**) 1D ^13^C CP spectra comparing rigid carbohydrates in WT and *ags1*Δ*ags2*Δ*ags3*Δ conidia at 0 h and 7 h prepared from 13C-labled conidia grown in unlabeled media. (**D**) Molar composition of rigid carbohydrates in WT (left column) and *ags1*Δ*ags2*Δ*ags3*Δ (right column) conidia harvested at 0 h (top row) and 7 h (bottom row).

Comparison of the three samples revealed extensive α-1,3-glucan turnover during germination. Modification of pre-existing cell wall components produced only a modest modification of resting conidium α-1,3-glucan during germination (**Figure 3A**; top), accounting for 17% of the rigid carbohydrates derived from pre-existing carbon pools (**Figure 3B**; top). In contrast, α-1,3-glucan accounted for 45-46% of the rigid cell wall carbohydrates in both the newly synthesized moleulces and the final, fully labeled samples (**Figure 3A, B**; middle and bottom). These results demonstrate that α-1,3-glucan neosynthesis is a major cell wall biosynthetic process during early germination and contributes substantially more than reutilization of pre-existing α-1,3-glucan.

Given the pronounced turnover and synthesis of α-1,3-glucan during germination, we next examined a triple α-1,3-glucan synthase deletion strain (*ags1*Δ*ags2*Δ*ags3*Δ) to investigate the consequences of α-1,3-glucan loss on cell wall organization. As expected, α-1,3-glucan signals were absent from the rigid cell wall fraction in both resting and germinating conidia, as indicated by the disappearance of the characteristic 101.0 ppm A1 peak (**Figure 3C**). The resulting mutant cell wall exhibited a simplified rigid core composed primarily of chitin and β-1,3-glucan, which accounted for approximately 31-36% and 64-69% of the rigid carbohydrate fraction, respectively, across all germination stages (**Figure 3D**). Compared with the wild type, loss of α-1,3-glucan triggered extensive compensatory remodeling of the cell wall, resulting in increased chitin accumulation in resting conidia (0 h) and concomitant increases in both chitin and β-1,3-glucan during germination (7 h), which likely compensate for the structural functions normally provided by α-1,3-glucan.

Notably, the α-1,3-glucan-deficient strain has been reported to exhibit a faster germination rate than the wildtype^[16, 27]^, consistent with a hypothesis that α-1,3-glucan-containing cell wall architectures contribute to constraining or regulating the timing of germination. It is likely that α-1,3-glucan, as a highly versatile cell wall component, plays important roles in shaping the polysaccharide remodeling pathways that accompany the transition from dormant conidia to actively growing germlings.

## DISCUSSION

This study employs high-resolution solid-state NMR spectroscopy and complementary ^13^C-labeling strategies to provide a comprehensive view of cell wall remodeling during *A. fumigatus* conidial germination. Tracking both pre-existing and newly synthesized carbon pools, while distinguishing rigid and mobile polysaccharide domains, revealed an extensive remodeling program that supports the transition from dormant conidia to actively growing germlings. A prominent feature of this process is the reduction of pre-existing conidial β-1,3-glucan during germination. The cell wall softening required for conidial swelling and germ tube emergence in filamentous fungi has often been attributed to the activity of specific glycosyl hydrolases that hydrolyze cell wall polysaccharides^[26, 28]^. The major β-1,3-glucan hydrolases belong to the GH16, GH55, and GH81 families in the CAZy database, comprising a total of 18 predicted β-1,3-glucan hydrolases. A fourth family, GH3, contains 10 glycoside hydrolases; however, none has been shown to possess true β-1,3-glucanase activity. Unexpectedly, deletion of the 28 β-1,3-glucanase genes did not impair the germination capacity of these mutants or result in complete or substantial hydrolysis of cell wall β-1,3-glucans. However, both endo- and exo-β-1,3-glucanase mutants exhibited characteristic conidial separation defects, forming linear chains of attached conidia consistent with the persistence of β-1,3-glucan-rich junction material^[25, 29]^. These results indicate that β-1,3-glucans play a major role during conidium formation and demonstrate that the glucan-rich septal material is structurally dynamic and that glucanase activity is particularly important during conidiation and conidial separation, a concept that is often overlooked in mycology^[30]^.

Moreover, degradation of β-1,3-glucan during germination does not exceed one-third of the cell wall β-1,3-glucan. This limited degradation suggests that β-1,3-glucan hydrolysis occurs only at specific sites, such as the apex of the emerging germ tube or sites of branch formation, rather than throughout the entire cell wall. These findings further suggest that inhibition of glucanase activity is unlikely to identify effective antifungal drug targets. In addition, GH132 transglycosidases, such as the SUN proteins, may play an unexpected role in cell wall remodeling during germination and have not yet been investigated^[31]^.

The glucan reduction is coupled with increased chitin and α-1,3-glucan that likely reinforce the expanding cell wall and maintain structural integrity. The continuous incorporation of neosynthesized α-1,3-glucan during germination also suggests that cell wall expansion is driven by an active remodeling program rather than by net wall degradation. Comparison of rigid and mobile polysaccharide fractions further revealed that this remodeling is spatially compartmentalized. The rigid cell wall undergoes extensive restructuring characterized by β-1,3-glucan turnover and α-1,3-glucan deposition, whereas the mobile fraction displays distinct dynamics. Galactomannan remains relatively stable throughout germination, while GAG is actively synthesized from both pre-existing carbon reserves and newly assimilated nutrients^[32–33]^. These observations demonstrate that individual cell wall polysaccharides are regulated through distinct biosynthetic and remodeling pathways during the transition from dormant conidia to actively growing germlings. In addition, intracellular metabolites such as trehalose, as well as other non-cell-wall components, may contribute to the biophysical properties of the cell wall and thereby influence its remodeling during germination^[34]^.

α-1,3-Glucans by themselves are not important for germination, even though this major polysaccharide, like chitin and β-1,3-glucans, is extensively remodeled during germination^[16]^. Instead, α-1,3-glucan and the extracellular polysaccharide GAG are important for the aggregation of germinating conidia and hyphae in *Aspergillus species*, such as *Aspergillus oryzae*^[35–36]^.

Earlier studies in *A. fumigatus* suggested that deletion of multiple chitinase genes has no effect on morphology despite a substantial reduction in chitinase activity^[37]^. Even though our study has shown that the chitin content of the cell wall changes during conidial germination, a direct involvement of chitinases in fungal growth remains under debate because of the lack of conclusive experimental evidence. Similar to β-1,3-glucanases, the functions of chitinases, based on the morphology of the corresponding mutants, have been proposed to be primarily involved in cell separation, sporulation, or chitin utilization during autolysis rather than in plasticizing the cell wall for germ tube emergence^[38–40]^.

As strongly suggested by the de novo synthesis data, the α-1,3-glucan synthase pathway appears to preferentially channel newly assimilated carbon into the rigid cell wall fraction during early germination, consistent with α-1,3-glucan serving as a primary investment polymer once nutrient uptake begins. The resulting changes in germination kinetics suggest that interactions among α-1,3-glucan, chitin, and β-1,3-glucan contribute to regulating the physical properties of the germinating cell wall^[41]^. Consistently, in more mature hyphae of *A. fumigatus*, α-1,3-glucan was found to associate with chitin and β-1,3-glucan, thereby reinforcing the rigid scaffold of the cell wall^[42–43]^. Meanwhile, the increased chitin content during germination may also have implications for susceptibility to chitin synthase inhibitors such as nikkomycin Z, although this possibility remains to be tested experimentally^[27]^.

This study confirms that glycosyl hydrolases, when considered individually, cannot control the initiation of germination. The functions of chitinases are similar to those of β-1,3-glucanases and appear to be more focused on cell separation than on plasticizing the cell wall. Although glycosyl hydrolases have been the subject of extensive research for decades, knowledge of their biological functions and their role in conidial germination remains limited, and interest in their study has diminished over the years. In addition, the linkages between cell wall polysaccharides have not been analyzed during germination, nor has the specific localization of the different hydrolases been investigated using super-resolution microscopy. Likewise, the enzymes responsible for forming linkages between polysaccharides, such as α-1,3-glucan and GAG or β-1,3-glucan and chitin, remain unidentified.

Our data are also consistent with previous studies confirming that germination is controlled by transcription factors regulating multiple pathways, such as the C2H2 zinc-finger transcription factors SltA, Ace2, and Swi5^[44–46]^. Key physiological processes in eukaryotic cells, including germination, are regulated by networks of protein kinase-based signaling pathways that control morphogenesis and polarized growth. This conclusion also suggests that it will be difficult to develop antifungal agents based on the identification of direct inhibitors targeting the activity of a single glycosyl hydrolase.

These biophysical results support a model in which *A. fumigatus* germination is governed by a robust and highly coordinated cell wall remodeling program that balances wall softening by hydrolases with structural reinforcement through biosynthetic pathways. Reduction of pre-existing β-1,3-glucan, together with de novo synthesis of α-1,3-glucan and chitin accumulation, enables swelling and polarized growth while preserving cell wall integrity. Remodeling is distributed across structurally distinct rigid and mobile carbohydrate domains and remains remarkably resilient to genetic or chemical perturbations through compensatory pathways.

A central conclusion of this study is that germination is not driven primarily by lysis of the pre-existing conidial wall. Instead, *de novo* polysaccharide synthesis is at least as important as, and may be more important than, putative polymer degradation during this developmental transition. This biosynthetic contribution has been largely overlooked because conventional biochemical approaches cannot readily distinguish inherited wall material from newly synthesized polymers. We directly differentiated the fate of pre-existing carbon from the incorporation of newly assimilated carbon into the cell wall, revealing substantial investment in new structural polysaccharides during early germination. These findings establish isotope-edited solid-state NMR as a powerful approach for quantitatively resolving polymer turnover and synthesis during fungal morphogenesis. They also show that the loss or modification of a single polysaccharide can be accompanied by compensatory changes in other wall components without an obvious or predictable pattern.

Future studies should characterize these reorganizations not only qualitatively but also quantitatively to define the subtle architectural changes and the role of polymer linkers that drive morphogenetic transitions, a concept often overlooked^[47]^. Cytochemical labeling and super-resolution imaging will be particularly valuable for defining where these modifications occur and for relating them to the regulatory pathways controlling germination^[48]^. Overall, these results establish the fungal cell wall as a flexible yet highly organized structure whose developmental remodeling depends on coordinated de novo synthesis and selective turnover rather than on generalized wall degradation alone.

## METHODS

### Preparing samples for solid-state NMR analysis

This study used four different *A. fumigatus* strains including the background wild-type strain (*akuB^ku80^*), α-1,3-glucan-deficient strain (*ags1*Δ*ags2*Δ*ags3*Δ), endo-β-1,3-glucanase-deficient strain (*engl1*Δ*eng2*Δ*eng3*Δ*eng4*Δ*eng5*Δ), and exo-β-1,3-glucanase-deficient strain (*exg8*Δ*exg9*Δ*exg7*Δ*exg5*Δ*exg6*Δ*exg10*Δ)^[7, 17, 49]^. The α-1,3-glucan-deficient strain, which lacks the respective polymer in the cell wall, was used as a control to examine the effects of α-1,3-glucan loss on germination. Likewise, the other strains lacked five endo-β-glucanases and seven exo-β-glucanases, which when present cleave β-1,3-glucan at internal sites and at free chain ends, respectively. These strains were included to determine how loss of β-1,3-glucan remodeling enzymes influences cell wall composition and structural changes during conidial germination.

Each strain was inoculated onto glucose minimal media (GMM) plates with 1.5% agar, 1% (w/v) ^12^C or ^13^C glucose (Fisher Scientific, Cambridge Isotope Laboratories respectively) as the sole carbon source, sodium nitrate salt solution (Cambridge Isotope Laboratories) as the sole nitrogen source and supplemented with trace elements (**Table S1**)^[50]^. The culture was allowed to grow for 5 days, incubated at 37°C and were harvested using 0.5% tween-20 solution and washed 3 times with nanopure water (Thermo Scientific) centrifuging at 8000 rpm for 3 minutes. Conidia thus harvested were subjected to growth in 100 mL of liquid GMM as follows: Labeled conidia (LC) in unlabeled liquid media (unLM), unlabeled conidia (unLC) in labeled liquid media (LM) and LC in LM. Each combination of conidia and media were shaken in an incubator at 37°C at 160 rpm for 0, 5 and 7 h, the sample then washed thrice, subjected to cell count in triplicate (**Table S2)** and packed into a 3.2 mm rotor for solid-state NMR measurement. At the same time, brightfield microscope images were captured on the Nikon C2^+^ Confocal Laser Scanning Microscope at the Centre for Advanced Microscopy at Michigan State University. The LCunLM combination is expected to show signals of ^13^C that were already in the conidia, while unLCLM is expected to show ^13^C from the media that were used by the fungus to neosynthesize cell wall components required by the fungus to germinate. LCLM would show a summation of both.

### 13C Solid-state NMR analysis of cell wall composition

The solid-state NMR experiments were carried out on a Bruker Avance Neo 800 MHz (18.8 Tesla) instrument equipped with a 3.2 mm HCN triple-resonance magic-angle spinning (MAS) probe at the Max T. Rogers NMR facility at Michigan State University. MAS frequency was set to 15 kHz for ^13^C detection, and the experiment temperature was held constant at 293 K. ^13^C chemical shifts were externally referenced to the tetramethylsilane (TMS) scale by calibrating the methyl signal of methionine-leucine-phenylalanine (MLF) powder at 14.0 ppm as a secondary reference. Chemical shift references and experimental parameters are listed in supplementary information. The structurally rigid components were investigated using CP and CORD sequences ^[51]^, which employed CP for initial magnetization transfer and a mixing time of 53 ms at 15 kHz MAS. Mobile regions were probed using refocused J-INADEQUATE spectra acquired under DP conditions with a 2s recycle delay^[52]^. The J-evolution segment comprised four 2.3 ms delays, tuned to optimize the intensity of carbohydrate moieties. Experimental parameters are listed in **Tables S3**, **4**.

Relative molar fractions of rigid and mobile carbohydrates were obtained using two complementary solid-state NMR experiments. Mobile carbohydrates were analyzed from 2D ^13^C-detected DP refocused J-INADEQUATE spectra, while rigid carbohydrates were evaluated using 2D ^13^C-CP CORD spectra. Peak intensities, areas, and volumes were measured in Bruker TopSpin 4.1.4. To reduce ambiguity from overlapping signals, quantification was limited to well-resolved resonances that could be confidently assigned to specific carbohydrate components. For each 2D dataset, cross-peak volumes assigned to the same polysaccharide were integrated and averaged across the selected resonances.

Spectra used for comparison were collected and processed using identical experimental and processing parameters. Integrated 2D cross-peak volumes were corrected for the number of scans prior to comparison. For each carbohydrate type, the normalized signal was further adjusted by the number of resonances contributing to that assignment, and the resulting values were expressed as a fraction of the total carbohydrate signal within either the rigid or mobile domain. Therefore, the reported compositions reflect relative molecular distributions rather than absolute carbohydrate concentrations, as summarized in **Tables S5**, **6**. Standard errors were calculated from the variation among integrated resonances used for each assignment, and combined uncertainties were determined by propagation of the individual errors.

## ACKNOWLEDGMENT

This study was primarily supported by National Institutes of Health (NIH) under award number R01AI173270 to T.W. The authors thank Melinda Frame for assistance with imaging.

## ABBREVIATIONS

AGS, α-glucan synthase; CAZy, Carbohydrate-Active enZYmes database; CORD, combined R2^n^-driven correlation spectroscopy; CP, cross polarization; DP, direct polarization; GAG, galactosaminogalactan; Gal*f*, galactofuranose; GalN, galactosamine; GH, glycoside hydrolase; GM, galactomannan; GPI, glycosylphosphatidylinositol; INADEQUATE, incredible natural abundance double quantum transfer experiment; LC, ^13^C-labeled conidia; LM, ^13^C-labeled medium; MAS, magic-angle spinning; NMR, nuclear magnetic resonance; TMS, tetramethylsilane; WT, wildtype.

