## Supplementary material for "Distinguishing Polymer Turnover from *De Novo* Biosynthesis Reveals Dynamic Cell Wall Remodeling During *Aspergillus fumigatus* Conidial Germination": SI 0723.pdf

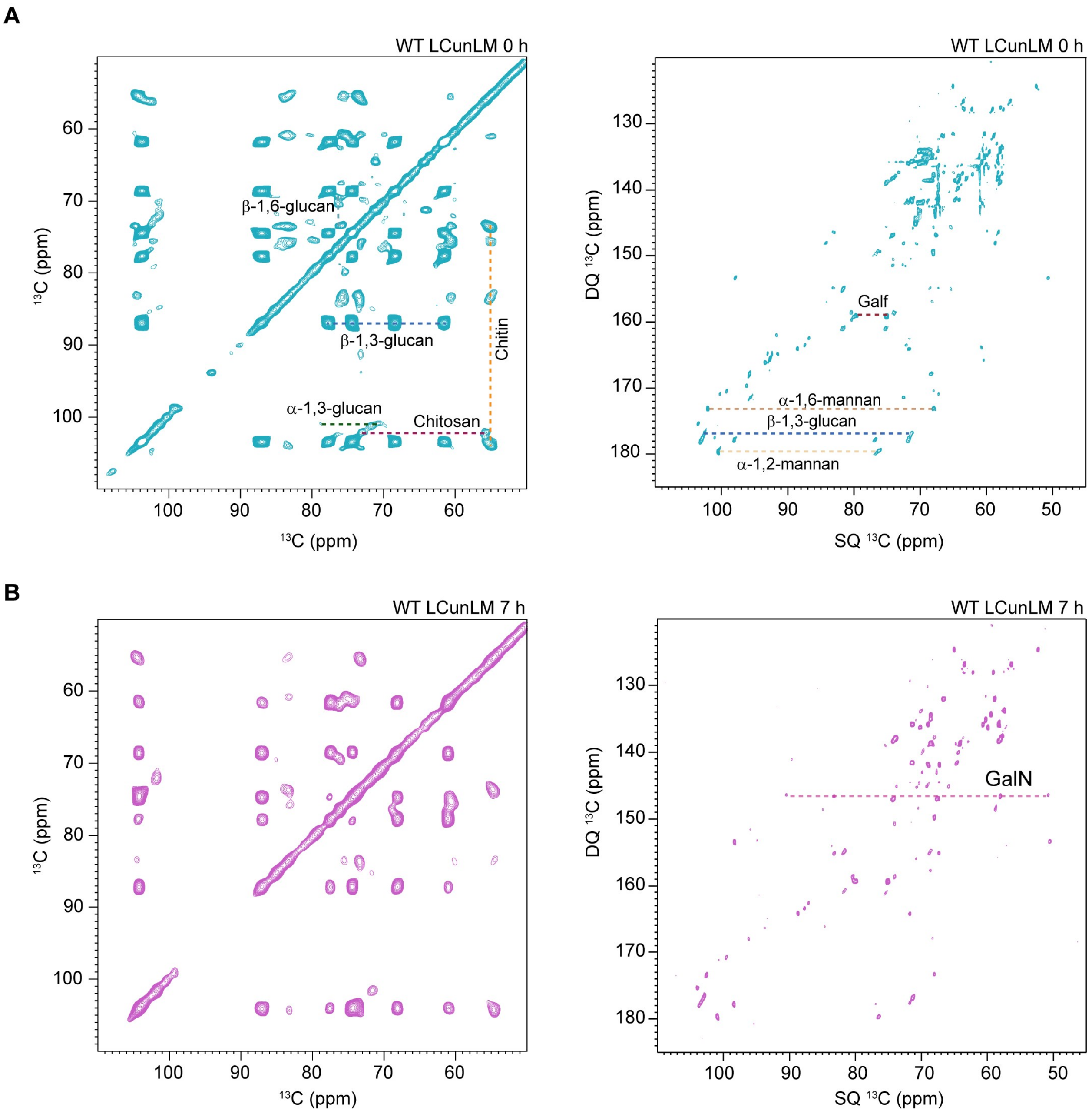


**Fig S1. Rigid and mobile carbohydrates in WT *A. fumigatus* conidia as it germinates.** (**A**) 2D $^{13}$C-$^{13}$C CORD (left) and $^{13}$C DP refocused J-INADEQUATE spectra (right) for resting conidia at 0 h. (**B**) 2D $^{13}$C-$^{13}$C CORD (left) and $^{13}$C DP refocused *J*-INADEQUATE spectra (right) for germinating conidia at 7 h.

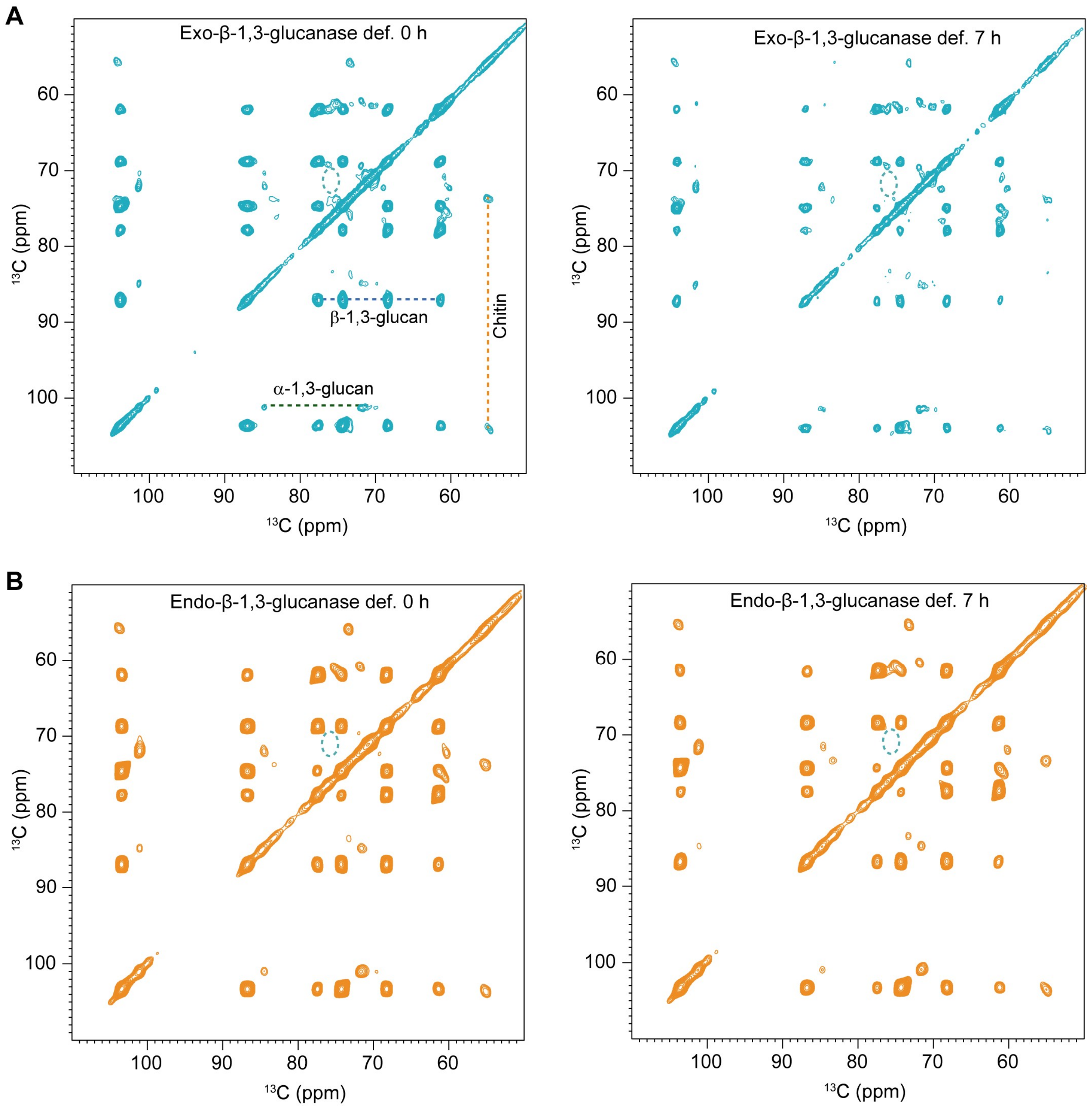


**Fig S2. Rigid carbohydrates in the conidial cell wall of mutant *A. fumigatus* as they germinate.** (**A**) 2D $^{13}$C-$^{13}$C CORD spectra for exo-β-1,3-glucanase-deficient strain at 0 h (left) and 7 h (right). (**B**) 2D $^{13}$C-$^{13}$C CORD spectra for endo-β-1,3-glucanase-deficient strain at 0 h (left) and 7 h (right). All 4 samples lack signals for β-1,6-glucan as shown by turquoise dashed lines.

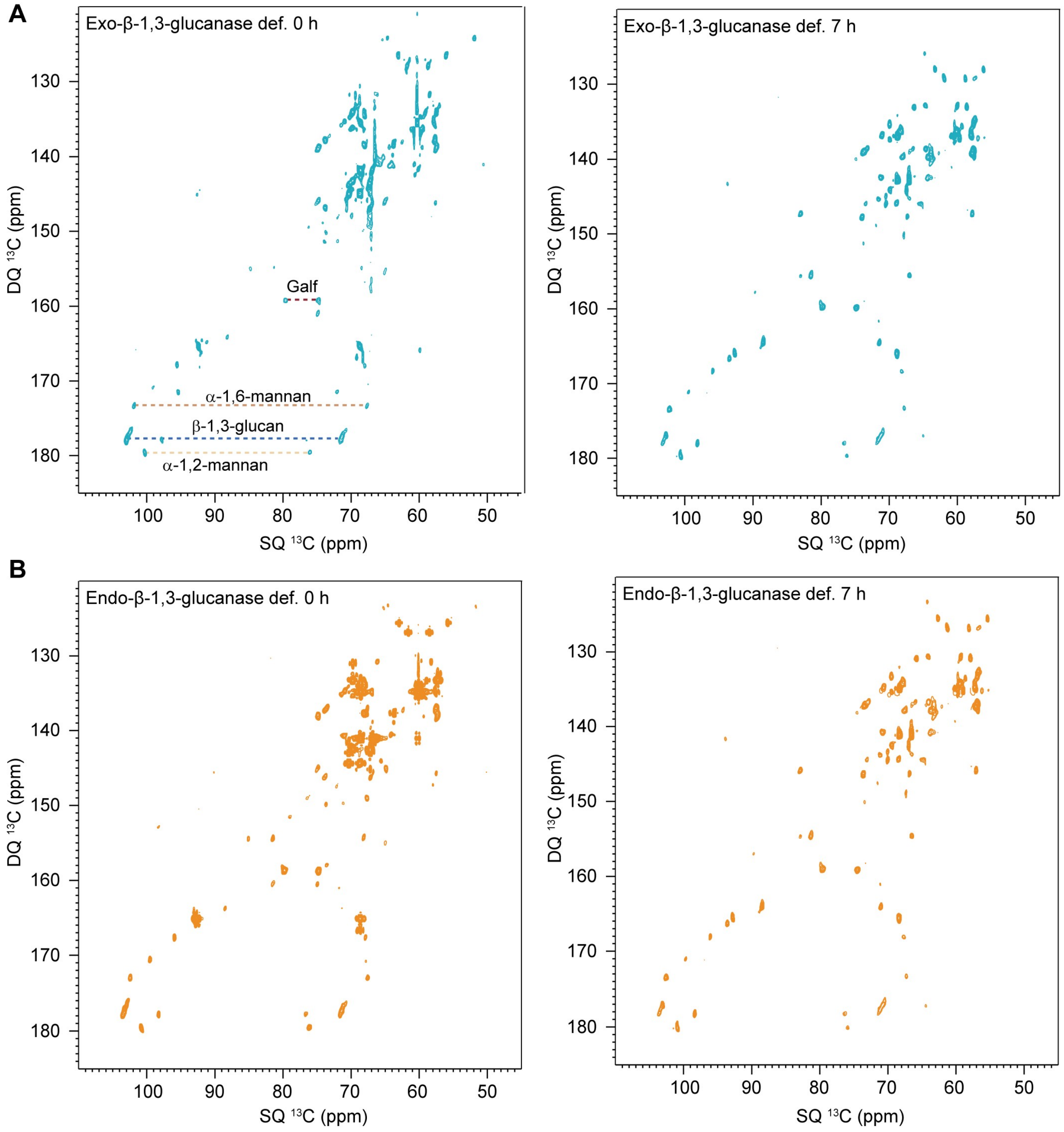


**Fig S3. Mobile cell wall carbohydrates in germinating mutant *A. fumigatus*.** (**A**) 2D $^{13}C$ DP refocused *J*- INADEQUATE spectra for exo-β-1,3-glucanase-def. strain at 0 h (left) and 7 h (right). (**B**) $^{13}C$ DP refocused *J*- INADEQUATE spectra for endo-β-1,3-glucanase-def. strain at 0 h (left) and 7 h (right).

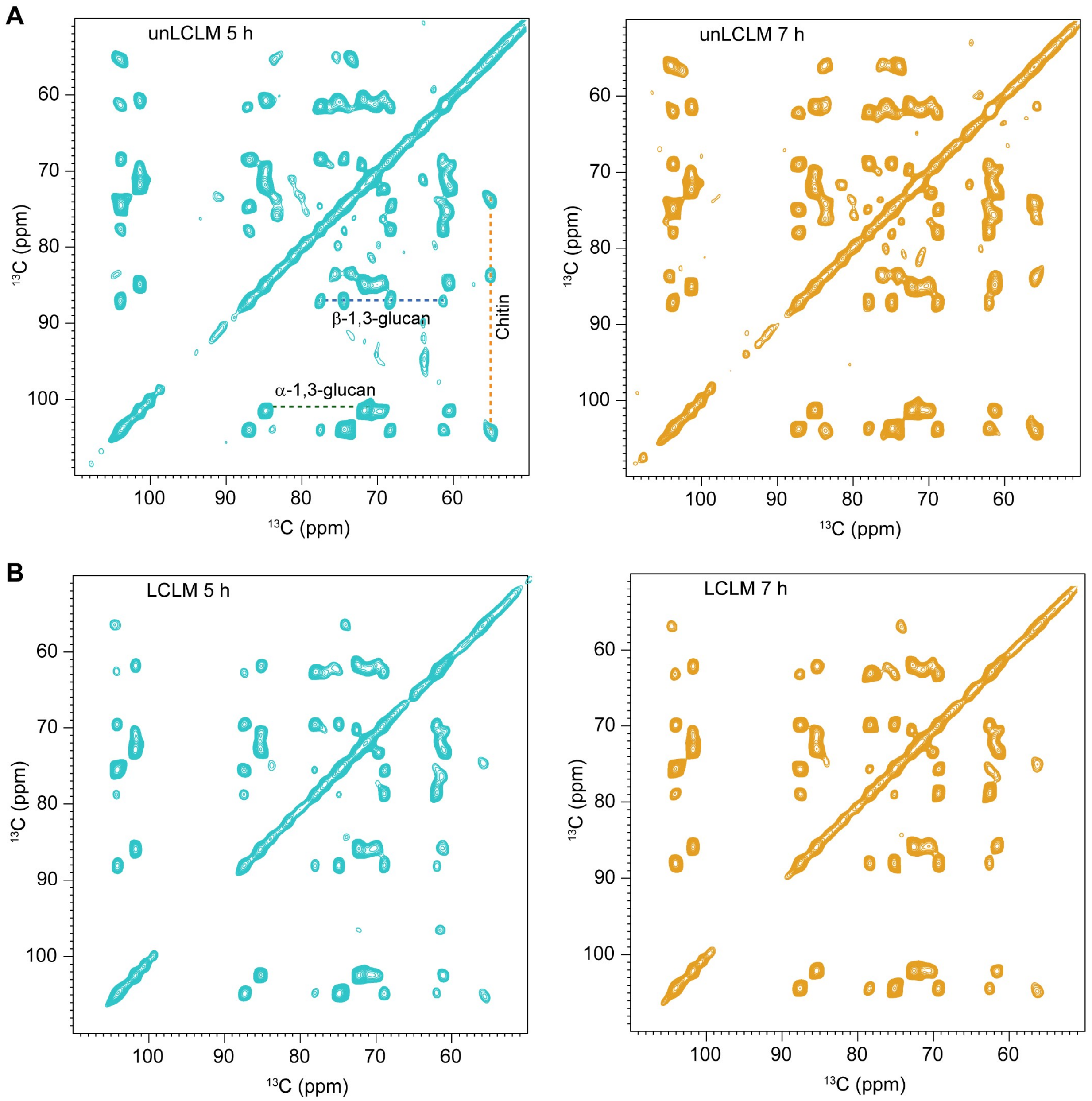


**Fig S4. Rigid carbohydrates in the conidial cell wall of germinating WT *A. fumigatus*.** (**A**) 2D $^{13}C$-$^{13}C$ CORD spectra for unLCLM samples at 5 h (left) and 7 h (right). (**B**) 2D $^{13}C$-$^{13}C$ CORD spectra for LCLM samples at 5 h (left) and 7 h (right). Both samples are lacking β-1,6 glucan signals.

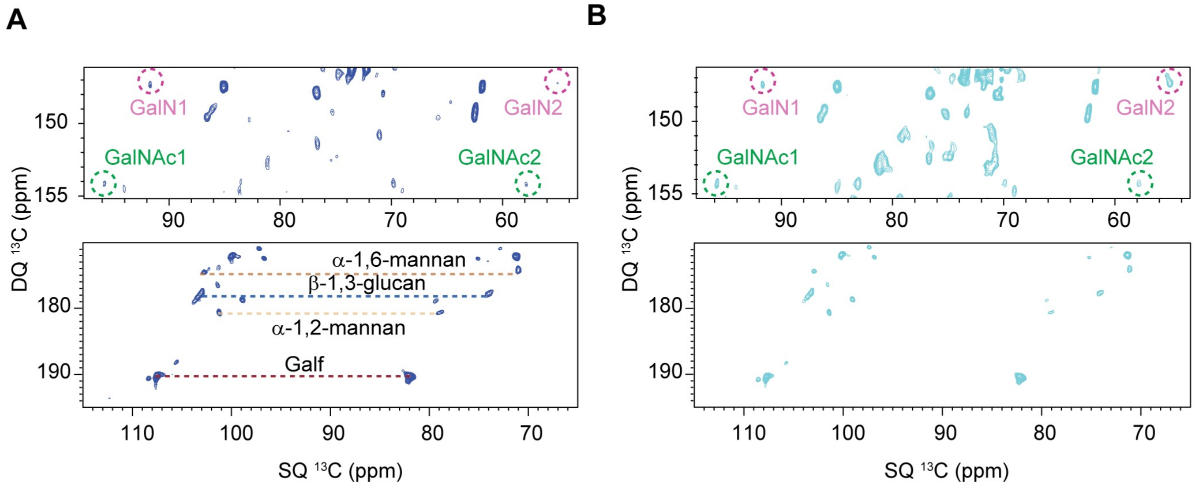


**Fig S4. Mobile carbohydrates in the conidial cell wall of germinating WT *A. fumigatus*. (A)** 2D $^{13}$C DP refocused *J*- INADEQUATE spectra for unLCLM samples at 5 h. **(B)** 2D $^{13}$C DP refocused *J*-INADEQUATE spectra for unLCLM samples at 7 h.

**Table S1**. **Recipe of mineral-based solid medium.** The pH is adjusted to 6.5 with $H_3PO_4$ or 0.25 M KOH. Each sample uses 100 mL of medium that contains 2g agar, 2 g of $^{13}C$-glucose and 5ml of sodium nitrate solution with 0.1ml of trace elements. The culture media and condition were adapted from a previously described protocol[1].

| | Reagent | For 1L |
|---|---|---|
| Trace elements | ZnSO4.7H2O | 22.0g |
| | H3BO3 | 11.0g |
| | MnCl2.4H2O | 5.0g |
| | FeSO4.7H2O | 5.0g |
| | CuSO4.5H2O | 1.6.0g |
| | CoCl2.6H2O | 1.6.0g |
| | (NH4)6Mo7O24.4H2O | 1.1.0g |
| | EDTA | 50.0g |
| Nitrate salt solution | NaNO3 | 300.0g |
| | KCl | 26.0g |
| | MgSO4.7H2O | 24.0g |

**Table S2. Summary of conidial counting using hemocytometer.** Conidial morphology was quantified by classifying counted conidia as resting, swollen, or germ tube-forming. For each replicate (n=3), the percentage of each morphological class was calculated relative to the average number of conidia counted, where the total conidia were ~$10^6$ per replicate. Data are presented as mean ± standard deviation from three independent replicates, rounded to the nearest whole number.

| | 0 h | | | 5 h | | | 7 h | | |
|---|---|---|---|---|---|---|---|---|---|
| sample | % resting conidia | % swollen | % germ tube | % resting conidia | % swollen | % germ tube | % resting conidia | % swollen | % germ tube |
| WT | 100 ± 0 | 0 | 0 | 20 ± 1 | 80 ± 3 | 0 | 2 ± 0 | 28 ± 1 | 70 ± 2 |
| Exo-β-1,3-glucanase def. | 100 ± 0 | 0 | 0 | 24 ±1 | 76 ± 2 | 0 | 2 ± 0 | 30 ± 0 | 68 ± 2 |
| Endo-β-1,3-glucanase def. | 100 ± 0 | 0 | 0 | 25 ± 0 | 55 ± 1 | 0 | 4 ± 0 | 32 ± 0 | 64 ± 1 |
| *ags1Δags2Δags3Δ* | 100 ± 0 | 0 | 0 | 12 ± 0 | 87 ± 2 | 0 | 3 ± 0 | 19 ± 0 | 78 ± 2 |

**Table S3. $^{13}$C chemical shifts of *A. fumigatus* cell wall and capsular polysaccharides in cell walls from $^{13}$C-based experiments**. The referencing scale is TMS scale. All chemical shifts are from room-temperature experiments.

| Carbohydrates | form | C1 | C2 | C3 | C4 | C5 | C6 | Reference |
|---|---|---|---|---|---|---|---|---|
| Rigid molecules | | | | | | | | |
| α-1,3-glucan (A) | a | 101.2 | 71.9 | 84.5 | 69.9 | 71.6 | 60.9 | Chakraborty *et al.* 2021[2] Shim *et al.* 2007[3] |
| | b | 101 | - | 84.7 | 69.2 | - | - | |
| β-1,3-glucan (B) | | 103.6 | 73.7 | 86.5 | 69.3 | 76.1 | 62.0 | |
| β-1,6-glucan (H) | | 102.9 | 74.1 | - | 70.2 | 76.4 | 69.2 | Lowman *et al.* 2011[4] |
| Chitin (Ch) | | 104.3 | 54.3 | 73.7 | 83.7 | 75.7 | 61.1 | Gautam *et al.*[5] |
| Chitosan (Cs) | | 102.4 | 55.6 | 73.1 | - | - | - | Cheng *et al.*[6] |
| Mobile molecules | | | | | | | | |
| β-1,3-glucan (B) | | 103.6 | 74.3 | 86.8 | 68.1 | 76.8 | 62.0 | Chakraborty *et al.* 2021[2] |
| α-1,2-Mannan ($Mn^{1,2}$) | | 101.3 | 78.6 | - | - | - | - | Chakraborty *et al.* 2021[2] |
| α-1,6-Mannan ($Mn^{1,6}$) | | 102.6 | 70.7 | - | - | - | - | |
| Gal*f* | | 107.4 | 81.5 | 77.2 | - | - | - | Chakraborty *et al.* 2021[2] |
| GalN | | 91.9 | 55.1 | - | - | - | - | Gautam *at al.*[5] |
| GalNAc | | 96.1 | 57.2 | - | - | - | - | |

**Table S4. Solid-state NMR experiments and parameters**. To be quantitative, direct pulse (DP) experiments with 35 s long recycling delay were used. cross polarization (CP), most rigid molecules. For 2D $^{13}C$-$^{13}C$ correlation experiments allowed to resolve rigid intramolecular peaks. 2D DQ-SQ, DP J-INADEQAUTE spectra were used to detect through-bond correlations. The experimental parameters include the $^{1}H$ Larmor frequency, total experiment time (t), recycle delay (d1), number of scans (NS), The number of points for the direct (td2) and indirect (td1) dimensions, the acquisition time of the direct dimension (aq2) and the evolution time of indirect dimension (aq1), spectral width (sw1 and sw2).

| Experiment | $B_0$ (T) | t (h) | d1 (s) | NS | td2 | td1 | aq2 (ms) | aq1 (ms) | sw2 (ppm) | sw1 ppm) | $\tau_{mix}$ (ms) | νMAS (kHz) |
|---|---|---|---|---|---|---|---|---|---|---|---|---|
| 1D $^{13}C$ CP | 18.8 | 0.5 | 1.8 | 1024 | 3600 | | 18 | | 496.8 | | | 15 |
| 2D $^{13}C$-$^{13}C$ with CORD | 18.8 | 11 | 2.0 | 32 | 2800 | 600 | 14 | 7.5 | 496.8 | 198.7 | 53 | 15 |
| 2D $^{13}C$-$^{13}C$ refocused DP J- INADEQUATE | 18.8 | 6 | 1.5 | 16 | 2600 | 1024 | 19 | 10 | 326.8 | 248.5 | | 15 |

**Table S5**. **Molar composition of rigid polysaccharides in *A. fumigatus* cell wall.** The numbers were calculated using the integrals of well-resolved cross peaks in 2D $^{13}C$-$^{13}C$ CORD spectra. The results were normalized by the number of scans.

| Sample | α-1,3-glucan ($A^a$) | α-1,3-glucan ($A^b$) | β-1,3-glucan (B) | Chitin (Ch) | β-1,6-glucan (H) | Chitosan (Cs) |
|---|---|---|---|---|---|---|
| WT LCunLM 0 h | 5 | - | 73 | 12 | 8 | 2 |
| WT LCunLM 5 h | 15 | 6 | 52 | 26 | - | - |
| WT LCunLM 7 h | 17 | 4 | 51 | 28 | - | - |
| WT unLCLM 5 h | 51 | - | 30 | 19 | - | - |
| WT unLCLM 7 h | 46 | - | 30 | 24 | - | - |
| WT LCLM 5 h | 43 | - | 42 | 15 | - | - |
| WT LCLM 7 h | 45 | - | 32 | 23 | - | - |
| Exo-β-1,3-glucanase def. 0 h | 9 | - | 81 | 10 | - | - |
| Exo-β-1,3-glucanase def. 7 h | 23 | - | 60 | 17 | - | - |
| Endo-β-1,3-glucanase def. 0 h | 11 | - | 75 | 15 | - | - |
| Endo-β-1,3-glucanase def. 7 h | 15 | - | 66 | 19 | - | - |
| *ags1*Δ*ags2*Δ*ags3*Δ 0 h | - | - | 69 | 31 | - | - |
| *ags1*Δ*ags2*Δ*ags3*Δ 7 h | - | - | 64 | 36 | - | - |

**Table S6**. **Molar composition of mobile polysaccharides in *A. fumigatus* conidial cell wall.** The numbers were calculated using the integrals of well-resolved cross peaks of β-1,3 glucan and chitin in $^{13}C$ DP *J*- INADEQUATE spectra. The results were normalized by the number of scans.

| Sample | β-1,3-glucan | $Mn^{1,2}$ | $Mn^{1,6}$ | Gal*f* | GalN | GalNAc |
|---|---|---|---|---|---|---|
| WT LCunLM 0 h | 40 | 19 | 15 | 26 | - | - |
| WT LCunLM 5 h | 32 | 21 | 19 | 28 | - | - |
| WT LCunLM 7 h | 30 | 20 | 18 | 27 | 5 | - |
| WT unLCLM 5 h | 31 | 19 | 16 | 21 | 7 | 6 |
| WT unLCLM 7 h | 30 | 18 | 17 | 19 | 10 | 6 |
| WT LCLM 5 h | 30 | 17 | 16 | 25 | 5 | 7 |
| WT LCLM 7 h | 29 | 16 | 17 | 20 | 8 | 10 |
| Exo-β-1,3-glucanase def. 0 h | 50 | 12 | 19 | 19 | - | - |
| Exo-β-1,3-glucanase def. 7 h | 39 | 21 | 23 | 17 | - | - |
| Endo-β-1,3-glucanase def. 0 h | 59 | 16 | 10 | 15 | - | - |
| Endo-β-1,3-glucanase def. 7 h | 44 | 20 | 19 | 17 | - | - |
| *ags1*Δ*ags2*Δ*ags3*Δ 0 h | 44 | 19 | 14 | 23 | - | - |
| *ags1*Δ*ags2*Δ *ags3*Δ 7 h | 24 | 19 | 16 | 41 | - | - |
